# Viral adaptations to alternative hosts in the honey bee pathogen *Paenibacillus larvae*

**DOI:** 10.1101/2024.09.01.610711

**Authors:** Keera Paull, Emma Spencer, Craig R. Miller, James T. Van Leuven

## Abstract

Bacteriophages (phages) that are intended to be used to treat bacterial infections are often improved using genetic engineering or experimental evolution. A protocol called “Appelmans” utilizes evolution in microtiter plates to promote the evolution of phages that can infect nonpermissive hosts. We tested a modification of the Appelmans protocol using the honey bee pathogen, *Paenibacillus larvae*. Three phages evolved together on four *P. larvae* strains following the standard Appelmans protocol and a modified version to ensure high phage diversity throughout ten rounds of passaging. The host range of 360 plaques were characterized and six new phage lysis patterns were identified. These new phage lysis patterns included plaque formation on previously nonpermissive, phage-resistant isolates that were used to identify phage types. The modified protocol did not drastically change the rate or number of new phage types observed but did prevent the phage population from being dominated by one phage that tended to rapidly raise in frequency. These findings showed how a minor modification of the Appelmans protocol influenced the development of phages for phage therapy. The method also provided improved phages for the treatment of bacterial infections in honey bees.

## INTRODUCTION

Honey bees are considered the most important and effective pollinators in agriculture [1,2]. The pollination performed by managed honey bee hives is essential to meet food production requirements for humans [3]. American Foulbrood Disease (AFB) decreases honey bee (*Apis mellifera*) populations, causing economic loss and potential food vulnerabilities. AFB is caused by the bacteria *Paenibacillus larvae* which infects and kills the honey bee larvae of a hive [4–7]. *P. larvae* is a spore forming, gram-positive, rod-shaped bacterium belonging to the class Bacilli. Honey bee larvae become infected with spores when they ingest contaminated food, followed by spore germination, proliferation of bacterial cells, and production of toxins that break down the epithelial lining of the larval digestive tract [8–10].

Treatment options for AFB are minimal and destructive, making it difficult to manage. Antibiotics are generally used only by commercial beekeepers in the US and are banned in most European countries. However, using antibiotics is undesirable in most cases because it contributes to the spread of antibiotic resistance [11–15] and necessitates additional testing before human consumption of treated bee products [16]. Currently, the only other treatment available for AFB is extremely destructive and focuses on curtailing spore spread by burning all affected equipment, hives, and bees. Bacteriophages (phages) might be a less destructive solution to treat AFB infections [17–19]. Using phages to kill pathogens—a process called phage therapy—can be safe and effective. Phages can target pathogenic bacteria without harming the natural microbiota. Studies using phage therapy methods to treat *P. larvae* infections have gained recent interest. Currently, there are around 50 characterized *P. larvae*-specific phages [20–25]. Further studies are needed to expand the *P. larvae*-specific phage bank and improve phage therapy efficacy against AFB. The specificity that makes phages ideal at minimizing off-target effects also makes a single phage unlikely to kill all of the strains of a pathogen in circulation. *P. larvae* diversity is currently organized into five phylogenetic groups – ERIC types I-V [26] and no one *P. larvae* phage can infect all types. ERIC types I and II are most commonly isolated, whereas types III and IV have not been isolated for years. Laboratory evolution is commonly used to improve phages. A method that has gained recent interest is called Appelmans protocol (Fig 1) [27]. The primary goal of this method is to adapt phages to new hosts, expanding the host range of a phage or phage cocktail. It has also been used to increase phage titers [28]. These evolutions are generally conducted on microtiter plates where a dilution series of phages are mixed with different hosts and incubated for a set time period (typically until the well appears clear, which indicates complete lysis). Cleared wells are combined and used to start a new round of evolution. Combining phages that grew on different hosts provides the opportunity for recombination to occur between phages, promoting evolution to new hosts. There is however the possibility for one phage to rapidly take over the entire phage population or for one phage in the cocktail to be outcompeted and dropout of the population. Both scenarios reduce the phage diversity. As the standard Appelmans protocol calls for combining all the cleared dilution wells for each host, if one phage grows very well, it can quickly become the numerically dominant phage in the pool. To avoid this, we tested a modification to the Appelmans protocol, where the original phage cocktail is added to the evolved cocktail between each round of evolution. Our phage cocktail consisted of three previously characterized phages of *P. larvae* (Table 1). These phages are all temperate and favor a lytic lifestyle [17,20,21]. Phage Fern and Scottie are known to infect all but one of the host species that we used (Table 2). Recent evidence from a study using *P. larvae* strain 3650 suggests that phage Fern may integrate into the host genome [29]. Phage-resistance via genetic mutations can also arise in *P. larvae* strain 3650 [29]. Here, we used three phage-resistant strains previously generated by our lab that were resistant to each of the three starting phages to help distinguish parental lysis patterns from phages with new lysis patterns that evolved during Appelmans.

**Table 1.** The three *P. larvae* phages used in this experiment. Average nucleotide identity (ANI) between the three phages likely impacts the frequency of phage recombination.

| Phage Strain | Accession Number | ANI to Fern | ANI to Scottie | ANI to Xenia |
| --- | --- | --- | --- | --- |
| **Fern** | NC_028851 | 100% | - | - |
| **Scottie** | NC_055030 | 50% | 100% | - |
| **Xenia** | NC_028837 | 75% | 50% | 100% |

**Table 2.**
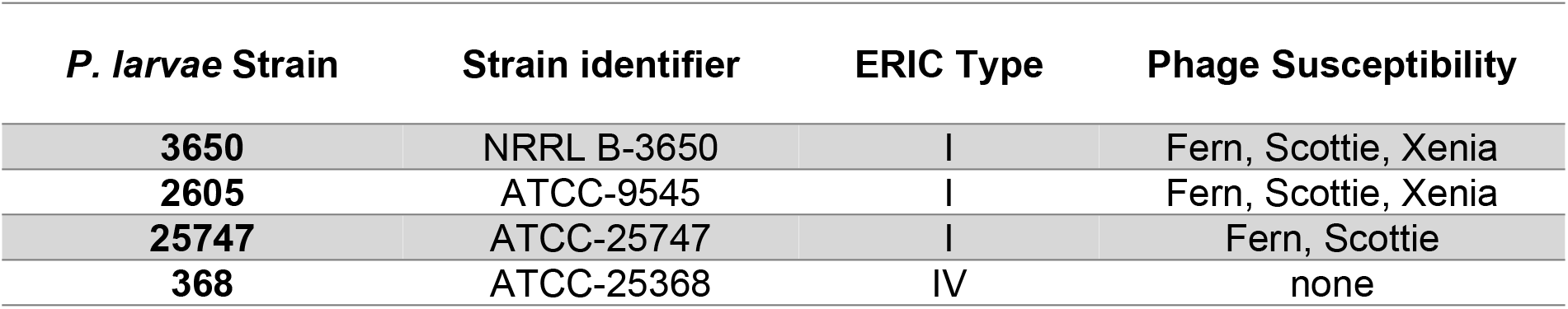
*P. larvae* strains used for Appelmans phage evolution.

**Figure 1.**
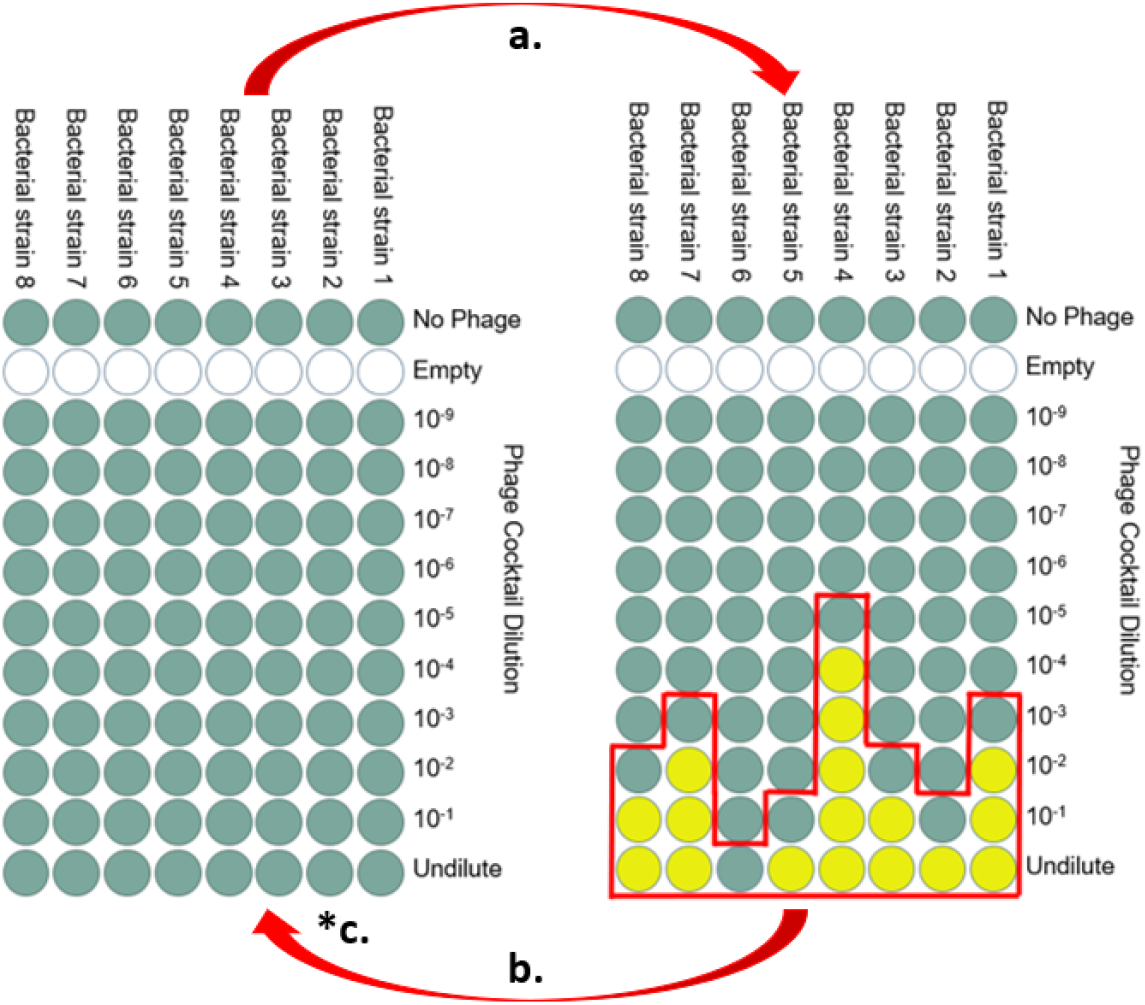
Standard Appelmans protocol to evolve phages on multiple bacterial strains. Each column of the 96-well microtiter plate contains a 200 μL culture of a single bacterial strain. Phages are added in ten-fold dilutions across the rows. (a) Overnight growth results in cleared wells (yellow) and turbid wells (green); (b) the next round of growth is started with a pool (red outline) of successful phages (the cleared wells) and the next turbid well from all columns; (*c) our modification: add 1:10 titer-based ratio of the starting phage cocktail to the pooled cocktail to start the next growth round.

## RESULTS

### Host range expansion

Over the course of ten rounds of passaging we randomly picked 18 plaques per time point for both the regular and modified protocols (18 plaques, 10 time points, 2 treatments = 360 total plaques). Only six new host range patterns were observed among these 360 plaques. On round three of passaging, we observed clearing of host 368 at the highest dilution of phage. This change was interpreted as host range expansion as the concentration of phage stock used to start each round of Appelmans was kept constant at 1E6 plaque forming units per mL (pfu/mL). However, no plaques were observed to form on host 368. Moreover, lysis of 368 was only observed in the well containing the most concentrated dilution of phage, suggesting that productive infection was not occurring, but rather a phenomenon called lysis from without. This indicates that if a phage had evolved to use 368 as a host, it did not improve over the ten rounds of evolution. Two phage isolates (“New 8” and “New 9”) created halos on host 368 (Table 3) but did not form plaques. Thus, a phage that can kill, likely by lysis from without, but not replicate on host 368 arose but stayed at relatively low frequency during the experiment. We observed true host range expansion of phage onto the phage-resistant isolates that we were using to identify phage types. A phage-resistant strain of 3650 was developed for each of the three phages used in the Appelmans protocol [29]. Inclusion of these phage resistant strains in the evolution experiment expands our ability to differentiate unique phage lysis patterns. One phage (“New 6”) could infect all the hosts except 368. The standard Appelmans protocol in fewer new patterns than our modified protocol. The modified protocol resulted in five of the six new phage lysis patterns, with two patterns unique to this experiment (Fig. 2). The standard protocol had one new lysis pattern, New 9, unique to the experiment, however it was one of the phages unable to amplify on 368 and XIIIβa, so it is only differentiated from phage Fern by its ability to kill but not form plaques on these two hosts.

**Table 3.**
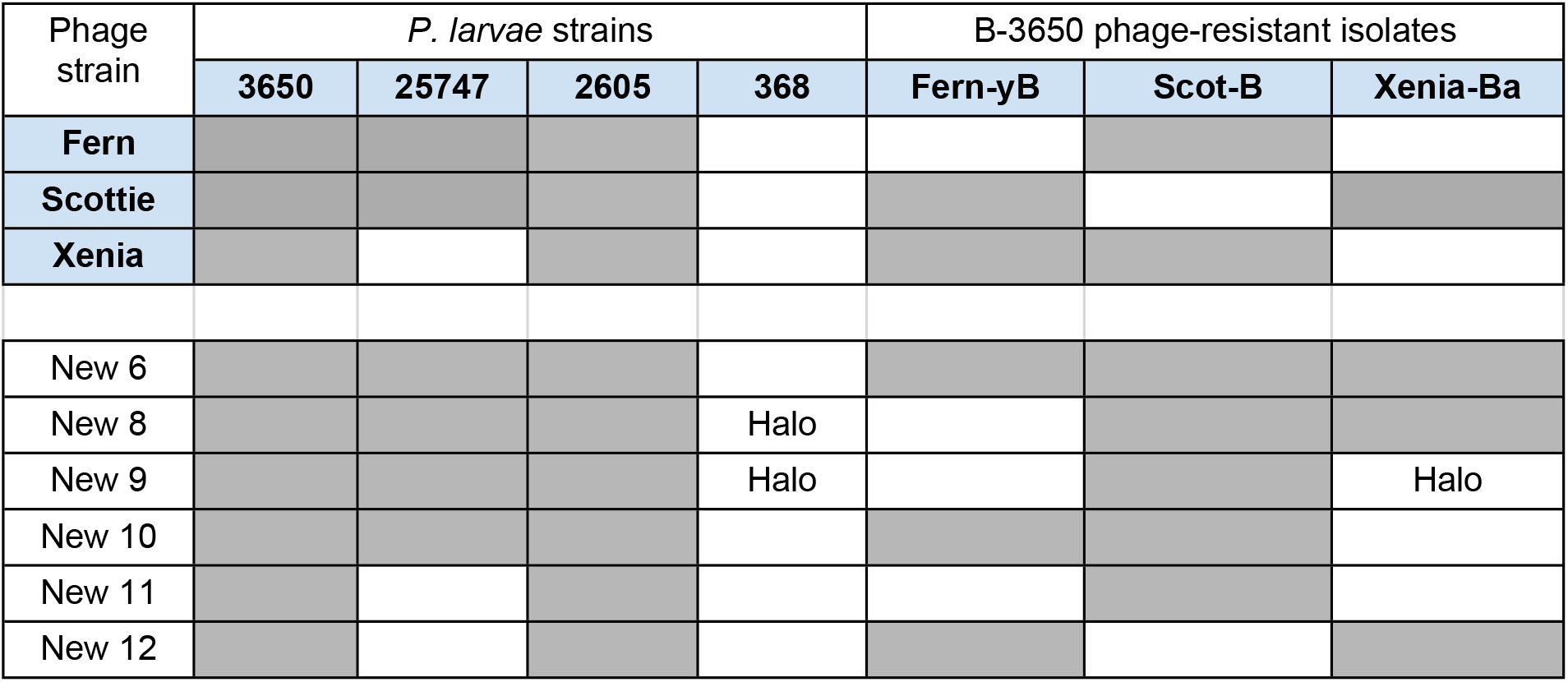
Host lysis matrix showing the three initial phages’ (Fern, Scottie, Xenia) ability to infect and kill the seven *P. larvae* strains used during evolution (top). Post evolution host lysis matrix showing the six new phage lysis patterns isolated during Appelmans (bottom 6 rows). Grey indicates host lysis, white indicates no host lysis, and “halo” indicates cell death without plaque formation.

**Figure 2.**
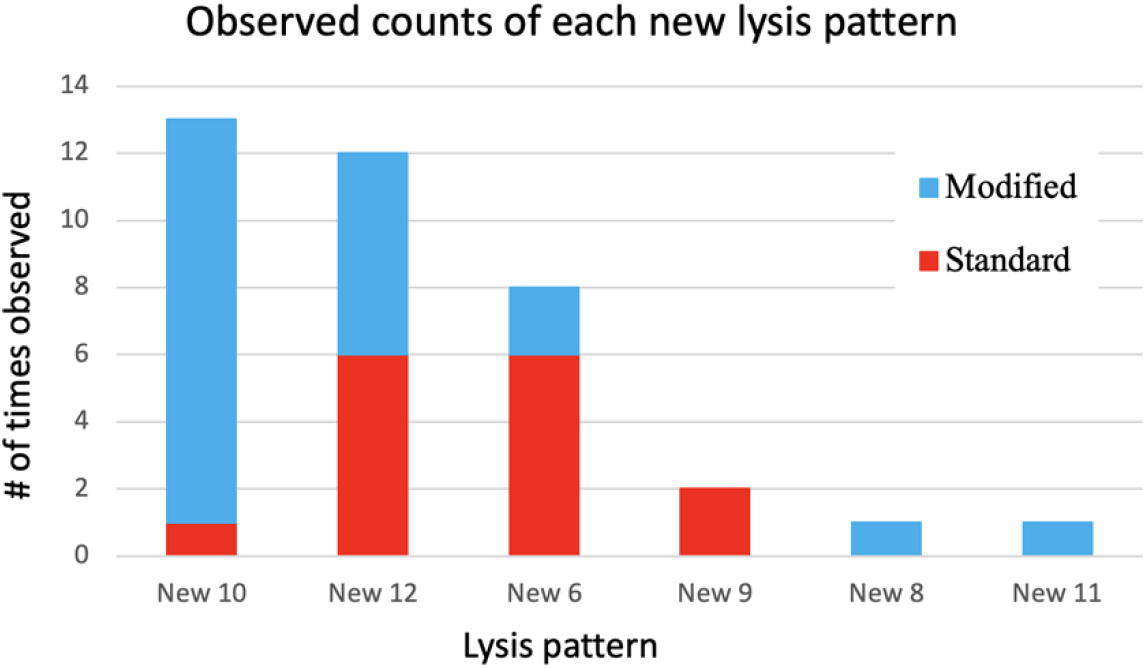
Number of times each new phage lysis pattern was isolated from the standard and modified protocols throughout 10 rounds of Appelmans evolution. These numbers were derived from the 18 randomly selected plaques from each round of evolution (360 total between both experiments).

### Timing and frequency of new phage types

For each of the ten rounds of passaging in the control and modified protocols, 18 plaques were isolated, amplified on 3650, and identified based on host-range patterns. New phage lysis patterns were isolated 22 times in the modified protocol experiment compared to the standard protocol, with 15 isolations (Fig 2). The first new phage lysis pattern was isolated in round two of the standard method and new lysis patterns were seen at a relatively consistent rate until the end of 10 rounds of our modified Appelmans protocol (Fig. 3). There were no major differences between the control and modified Appelmans protocols in terms of the rate that new types of phages arose. The new phage types made up a relatively higher proportion of the population in the modified protocol experiment compared to the standard protocol (Fig 4). The modified protocol also maintained higher phage diversity.

**Figure 3.**
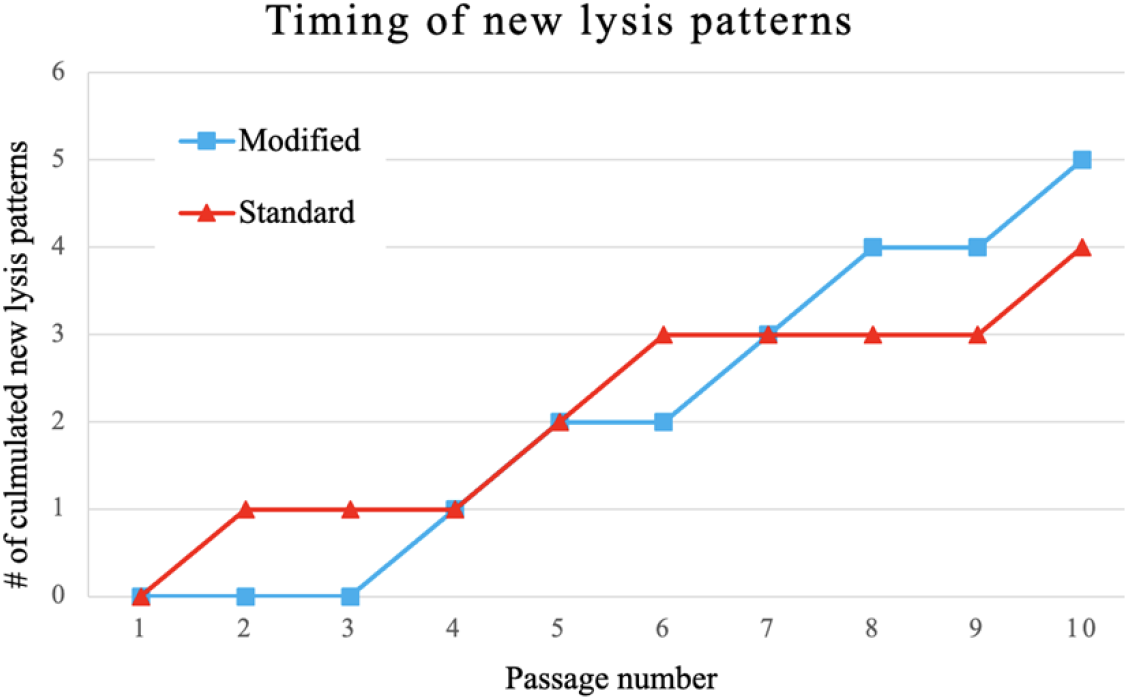
Accumulation of new phage lysis patterns from 360 picked plaques. Host range patterns for each plaque identified by plating on the seven hosts in Table 3.

**Figure 4.**
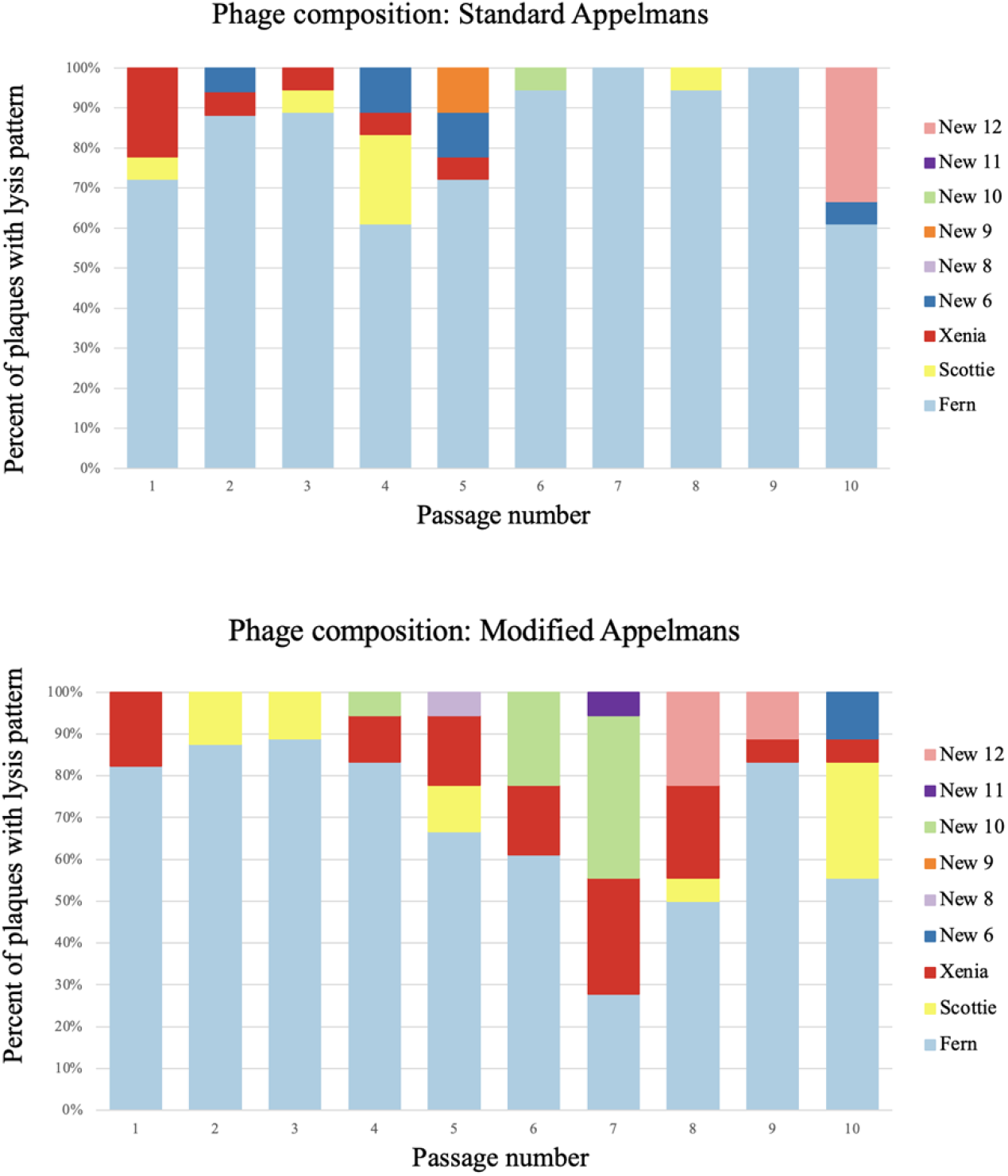
Phage lysis pattern composition for each round in the standard (top panel) and modified protocol (bottom panel) experiments. Each bar represents the composition of 18 plaques that match phages with the lysis patterns shown in the key.

Whereas phage Xenia was completely lost in the standard protocol experiment, it was present throughout the rounds of evolution in the modified protocol experiment. In both protocols, phage Fern was numerically dominant in the phage population. This highlights one concern of the standard Appelmans protocol. One should pay attention to the relative growth rates and number of permissive hosts for each phage going into Appelmans. In our case, we started with an equal mix of phage Fern, Scottie, and Xenia. It would be interesting to see the outcome if this starting mix would have been included a much lower titer of Fern.

### Dynamic phage population

The Appelmans protocol calls for pooling of all the cleared wells (indicative of lysis) plus the next turbid well (bacteria still growing with the more dilute phage cocktail present) for each host. We measured the phage titer of these lysates in each round and then always started the next round with 1E6 pfu/mL. The titer of the phage population on host 3650 did not change drastically over the ten rounds of evolution, always staying between 5E6 and 5E7 pfu/mL. However, we noticed drastic differences between hosts in the dilution of phage that resulted in cleared wells. Comparing the minimum number of phages required to clear a well of each host across the experiment shows these dynamics (Fig. 5). For susceptible hosts, it generally took around one phage to result in a cleared well. That is to say that wells containing susceptible hosts and that we expect to have at least some phage in them, based on the starting titer of 1E6, generally become clear. A decrease in the number of phages it took to clear a well indicated that the phage cocktail increased in efficiency of killing previously unsuceptible *P. larvae* strains. This can be seen clearly in the strain Fγb at round four for the standard and modified experiments (Fig 5). For the other hosts, we observed stochastic variation around one, indicating that the efficiency of killing was not dramatically changing for most hosts. The fluctuations are an unexpected result could indicate fluctuations in populations of phages specific to each of the hosts or that small numbers of phages are unreliably measured at the highest dilutions. Either way, we frequently observe wells clearing that were not expected to have phages in them based on the 1E6 starting titer and that this is not consistent across all the hosts. We may see an increase in the highest dilution cleared well for one host, but not the others.

**Figure 5.**
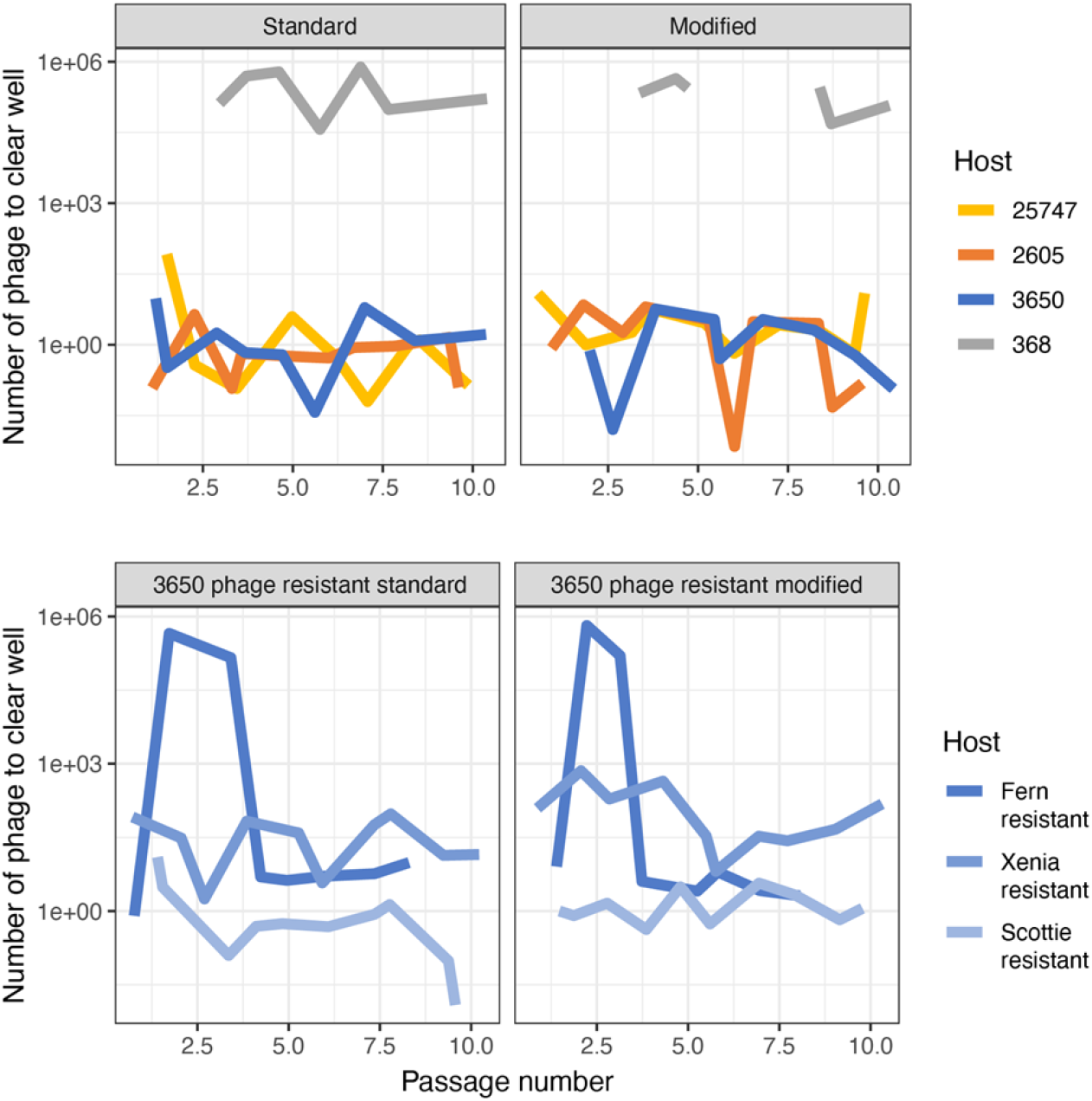
The initial number of phages (1E6 pfu/mL) divided by the phage dilution of the last cleared well for each *P. larvae* strain after every round of growth for both the standard and modified protocols. (a) naïve *P. larvae* strains; (b) phage-resistant *P. larvae* strains.

## DISCUSSION

As phage therapies for American Foulbrood disease are developed in honey bees, an improved understanding of the bacteriophages and how they evolve with their hosts is needed. The complications of phage-resistance and narrow host range are among the issues that would benefit from more research. Appelmans protocol has become a popular method of co-culture that can be used to improve many characteristics of a phage cocktail. As the use of Appelmans expands, several interesting complexities have arisen and more detailed studies of the population dynamics in these experiments will lead to an improved understanding of the method and of phage-host evolution in general [31]. We compared two strategies of evolution to determine if improvements could be made that accelerate the development of *P. larvae* phages. Our result show no substantial increase in the number of phages with new host range profiles when we modify the standard Appelmans protocol with the addition of phage each round of passage. This result may have been limited by the small total number of new host range profiles that we observed (only 6 from 360 tested plaques). Most reports using Appelmans (or some variation of it) result in a phage cocktail that is able to infect more hosts than the starting phage mix [27,32–34] but often the exact phage composition and genetic underpinnings of host expansion are not resolved in these studies. When these details are determined the results are often surprising. In a study of *Acinetobacter baumannii*, the host range of 56 phage isolates were tested after Appelmans and 12 isolates with new host range patterns were identified. Our rate (6 of 360 plaques) was substantial lower than this. While several of the *A. baumannii* isolates had gained the ability to grow on a novel host, some actually lost the ability to grown on hosts that they could originally replicate on. Sequencing of these phages showed frequent recombination and the presence of phages not in the starting phage cocktail. Induced prophages contributed to the host range expansion of the phage cocktail. Recombinants of the original phages did not have expanded host range. Temperate phage induction during Applemans has been reported in other studies [34] and highlights an unexpected benefit—or challenge—of using evolution on hosts to improve phages. In the *A. baumannii* study, host expansion was slower than in other applications of Appelmans, with around 70% of the hosts becoming susceptible to the phage cocktail at the end of the experiment. Studies on *Pseudomonas aeruginosa* found 100% of the hosts to be susceptible after a similar experiment [27,33,34]. Since we only had seven hosts, three of which were mutants of the original strains, it is hard to say exactly what this percentage was in our experiment. All seven hosts were killed by the phage cocktail, but 368 never supported phage replication. The modified Appelmans protocol did produce two unique phage lysis patterns, suggesting that perhaps this modification provides some benefit to produce novel phages.

A main difference that we observed between the standard and modified Appelmans protocols was the maintenance of a higher amount of phage diversity in the modified protocol. Therefore, addition of the starting phage cocktail to the pool of successful phages each round is a simple step that can benefit the evolution of phages in a cocktail. In our set of phages and hosts, one quickly rose in frequency. In other systems, this may not be the case and the improvement of our modified protocol may not be as necessary.

One key to successful phage therapy is the ability for the phages to infect the target bacteria, replicate, and lyse the host, thus killing it. This lytic infection cycle ensures host death along with amplification of the phages. Expansion of our phage cocktail’s host range to a nonpermissive *P. larvae* strain with Appelmans evolution was expected to derive phages capable of this lytic infection cycle on new hosts. Due to the clearing observed on host 368 in the undilute wells during Appelmans evolution and partial clearings seen on host range assay plates, new phage lysis patterns New 8 and New 9 were initially described as being able to infect and kill *P. larvae* strain 368, a previously nonpermissive host to phage infection. However, upon further testing no phage replication and amplification was observed on this host. There was clearly death of this *P. larvae* strain though. We predict that there is some mechanism of induced cell death occurring that is triggered by the presence of phages. Situations where a host cell kills itself or stops cellular functions to avoid phage infection is called abortive infection [11]. Because the killing of 368 did not arise until round two of evolution, we hypothesize that phages evolved to infected 368, but the infection is thwarted at the phage replication step by some phage-defense mechanism. Further genetic and molecular analysis of *P. larvae* strain 368 is needed to accurately describe why bacteria cells are dying in the presence of phages rather than directly due to phage infection.

Of the seven *P. larvae* strains, only one strain, 368, was nonpermissive to all three phages, Fern, Scottie, and Xenia, initially. Therefore, host range expansion of the phage cocktail with 10 rounds of a modified Appelmans protocol was unfortunately unsuccessful in this case. Increasing the number of evolution rounds may provide more time and opportunity for novel phage mutations to occur and expand the lytic life cycle to nonpermissive *P. larvae* strains.

*P. larvae* strains permissive to phage infections will likely evolve to develop resistance to phage infection just as they do to antibiotics [6, 17]. To further characterize these selective pressures between phages and *P. larvae* specifically, developed phage-resistant *P. larvae* strains were used during Appelmans evolution. New phage lysis pattern New 6 is capable of infecting and killing all three of the phage-resistance *P. larvae* strains, a host range pattern not seen in any of the starting phages. These results demonstrate that the inevitable host resistance of *P. larvae* can be overcome by phages evolved together in a cocktail. Therefore, using this evolved phage cocktail will likely benefit the success of phage therapy when treating persistent American Foulbrood infections. Future work will seek to identify the genetic determinants of these evolved phages to identify genes involved in host range expansion.

Part of making phage therapy a successful treatment is reducing the amount of time it takes to produce an effective phage cocktail, whether that includes evolution or not. If evolution is needed, our Appelmans protocol can produce new phage lysis patterns in as few as two rounds. Adding the ancestral recombination phage cocktail also showed an increase in individual new phage lysis pattern isolated likely due to more genetic recombination between phages. This addition also produced a diverse phage cocktail with a more proportional composition between the phage lysis patterns observed. This benefits phage therapy applications because a more diverse phage cocktail will likely be able to avoid total host resistance.

This research will benefit the pursuit of using phages to treat American Foulbrood disease in honey bees along with defining phage evolution protocols that can develop a more diverse, and therefore effective, phage cocktail.

## METHODS

### Bacterial and phage strains, maintenance, and culturing

*Paenibacillus* phages Fern, Scottie, and Zenia and *P. larvae* strains NRRL B-3650 and ATCC-25368 were acquired from Dr. Penny S. Amy at the University of Nevada Las Vegas. Strains ATCC-9545 (B-2605) and ATCC-25747 were ordered directly from the American Type Culture Collection. These bacterials strains will henceforth be referred to as 3650, 368, 2605, and 25747, respectively. The phage resistant isolates used in this study were generated in our lab [29] and will be referred to as Fγb (resistant to Fern), Scotβ (resistant to Scottie), and XIIIβa (resistant to Xenia).

Frozen stocks were made from overnight cultures grown to 0.7 OD. 1.5 mL of the culture was aliquoted into 2 mL freezer tubes, spun down at 11000 xg for 10 minutes, and the supernatant was discarded. 750 μL of fresh media and 30% glycerol was added to each tube, vortexed thoroughly, and then flash frozen in liquid nitrogen. All frozen stocks were stored at - 80°C and used to start experiments.

All *P. larvae* strains were continuously grown in 1 mg/L of thiamine hydrochloride Brain Heart Infusion broth at 37°C in a shaker incubator at 200 rpm for 24-48 hours [30]. Each shaking broth culture was started from an individual colony picked from streaks made on 1 mg/L of thiamine hydrochloride BHI agar plates. Plates were incubated inverted at 37°C with 5% CO_2_ for 24-48 hours. Strains were grown with 5% CO_2_.

### Plaque picking and double isolation

For phage isolation, one clean plaque was picked using a cut pipette tip and put in 750 μL of mBHI broth, 10% v/v chloroform was added, vortexed, and then spun down at 20000 x g for 5 minutes. 500 μL of the supernatant was put in a new tube and titered on host strain 3650. Phages were stored at 4°C and used as working stocks.

### Bacteriophage titering

Soft mBHI top agar overlays were used to determine phage concentration. 100 μL of *P. larvae* were mixed with varying amounts of a phage stock dilution in 3 mL of warm (50°C) mBHI, 1% agar. The mix was briefly vortexed then poured on top of a BHI agar plate. Plates were incubated upright at 37°C with 5% CO_2_ for 24-48 hours. Plates ranging from 50-250 plaques were counted and then averaged. Phage stock titers were calculated based on the average plaque count and dilution plated.

### Bacteriophage host range assays: spot plating

A grid pattern with nine boxes was draw on the bottom of BHI agar plates, one for each phage and one control. Soft mBHI top agar with 100 μL of a *P. larvae* strain was poured on the plate and let sit for an hour. 10 μL of undiluted to 1.0E-2 dilution (depending on titer) of each phage stock was pipetted into its labeled gridded box on the top agar layer and 10 μL of mBHI was pipetted into the control box. Plates sat for another hour to prevent unwanted spreading or mixing of phage and then incubated upright at 37°C with 5% CO_2_ for 24-48 hours. This was repeated for all the *P. larvae* strains. The next day lysis patterns of each phage were observed and recorded.

### Ancestral bacteriophage recombination mix culturing

To maintain and increase diversity in the phage cocktail, we implemented a modification to the standard Appelmans protocol that involved adding in a mix of the original phages before each new round of evolution. To obtain this mix, we cross streaked phages Fern, Scottie, and Xenia on one plate and obtained phages from the center of the plate where they were co-localized. Briefly, a 100 μL 3650 soft overlay top agar plate was prepared and let sit for an hour at 37°C. 10 μL of each phage stock was pipetted on the edges of the plate at equal distances from each other. Using a sterile loop, the drops were streaked straight across the plate crossing in the center. The plate was incubated upright at 37°C with 5% CO_2_ for 24 hours. The next day, a punch from the center clearing was collected in a 5 mL tube with 2 ml mBHI, mashed with a sterile tube mortar, vortexed, chloroformed, and centrifuged at 20000 x g for 10 minutes. The supernatant was collected and titered on 3650. This became the “ancestor recombination mix” fridge stock. This was done to allow growth and recombination between the three phage strains and later be added into each round of the Appelmans recombination series.

### Appelmans and modified Appelmans protocol

The general methods outlined in Burrows et al. (2019) [27] were followed with some minor modifications. 3 mL BHI broth cultures started from a colony were used to start overnight cultures of all *P. larvae* strains, 3650, 25747, 368, 2605, Fγb, Scotβ, and XIIIβa. The next day bacterial cultures were normalized to 0.3 OD at 600nm. For ease of pipetting, 1.0E-1 dilutions were made of the normalized hosts. The starting phage cocktail, equally comprised of phages Fern, Scottie, and Xenia, had a combined total titer of 1.0E6. The phage cocktail and all ten-fold dilutions of the cocktail (1.0E-1 to 1.0E-9) were in mBHI broth.

A 96-well microtiter plate was used for each round of evolution. Each column (12 wells) was used for one *P. larvae* strain and each row, 8 wells, was used for a dilution of the phage cocktail or control, see Figure 3. Row 1 control was media along with the *P. larvae* to monitor bacterial growth without phage infection. Row 2 control was just media (no bacteria or phage) to monitor media contamination. 100 μL of double strength mBHI broth was added to all 96 wells. 100 μL of mBHI was then added to only the row 2 control. For each column, 10 μL of a 1.0E-1 dilution of the normalized *P. larvae* strain was added to every well but the second (row 2 control). 100 μL of the corresponding phage cocktail dilution was added to each row skipping control rows 1 and 2. The most dilute phage cocktail (1.0E-9) wells were closest to the control rows, whereas the most concentrated phage cocktail (undilute) wells were in the row farthest from the controls (bottom row). For the first round of Appelmans, this set up was done twice to start both the control and recombination series.

The microtiter plates were covered with a Breathe-Easy membrane to allow gas exchange while preventing phage and bacterial contamination during growth. Plates were incubated overnight (18-24 hours) at 37°C in a shaker incubator (120 rpm).

The next day, photos were taken of each plate and the cleared wells (lysis) were recorded. 100 μL from each of the cleared wells from all hosts along with the next turbid well (next dilution) were pooled together. This was done separately for each series. The pooled lysate was then chloroformed (10% v/v), vortexed, and spun down at 22000 xg for 10 minutes. The supernatant (normally around 3 mLs) was put into a new tube and stored in the fridge for the next round. 1 mL freezer stocks were made from each round’s pooled lysate as well.

The pooled lysate from each round was tittered on *P. larvae* strain 3650 to determine the number of phages that will start the next round. For every round besides the first, the recombination series includes the modification where the ancestral recombination phage cocktail was added to the rounds pooled lysate based on a 1:10 titer ratio. Based on the recombination series pooled lysates titer and the known titer of the ancestral recombination phage cocktail, the 1:10 mix was made to start the next round (1 part ancestral recombination phage cocktail titer to 10 parts recombination series pooled lysate titer). The control series did not include this step, however it was also tittered after each round for later data analysis.

The set up for the next round of evolution is the same as the first round, but the phage cocktail and dilutions are made from the previous rounds pooled lysate cocktail and pooled lysate cocktail plus the ancestral recombination cocktail for the control and recombination series respectfully. This was repeated for ten rounds of growth.

### Appelmans pooled lysates host range determination and isolation

All 18 isolated phages from each round were picked from the 3650 toothpick plate for double isolation using the method described above. Once double isolated stocks were made for all 360 picked plaques (180 control series + 180 recombination series), host range assays using the spot plating method were conducted on all 8 hosts (4 naïve *P. larvae* strains, 3 phage resistant *P. larvae* strains, and out outgroup *P. alvei* strain B-383). There was never any lysis on the *P. alvei*, so its results were not included in data analysis.

Host lysis patterns were characterized and analyzed to predict which phage was present based on the ancestral host lysis patterns. If the observed host lysis pattern did not match any of the ancestral, it was named “new phage lysis pattern x” where x is any number. Spot plating host range assays were repeated to confirm predicted phage lysis patterns.

## ACKNOWLEDGEMENTS

We thank Drs. Penny Amy, Kurt Regner, Philippos Tsourkas, and Diane Yost for assistance in obtaining the *P. larvae* phages. Thank you to LuAnn Scott for laboratory assistance and training, Dr. Holly Wichman for experimental resources and comments, and Dr. Tracey Lee Peters for her helpful comments on this study, and manuscript edits.

